# Short-Term Warming Induces Cyanobacterial Blooms and Antibiotic Resistance in a Freshwater Lake as Revealed by Metagenomics Analysis

**DOI:** 10.1101/2024.07.08.602497

**Authors:** Bharat Manna, Emma Jay, Wensi Zhang, Xueyang Zhou, Boyu Lyu, Gevargis Muramthookil Thomas, Naresh Singhal

## Abstract

Climate change poses significant risks to freshwater ecosystems, potentially exacerbating harmful cyanobacterial blooms and antibiotic resistance. We investigated these dual threats in Cosseys Reservoir, New Zealand, by simulating short-term warming scenarios and assessing the role of oxidative stress. Microcosms were subjected to Base (22°C), Normal (24°C), and projected Future (27°C) temperatures, with additional treatments including reactive oxygen species (ROS). Metagenomic analysis revealed substantial restructuring of microbial communities under warming conditions. Cyanobacterial abundance increased from 6.11% (initial) to 20.53% at 24°C and 10.66% at 27°C. Notably, ROS addition mimicked the effects of temperature increase on cyanobacterial proliferation. Toxin-producing families, including *Microcystaceae* and *Nostocaceae*, proliferated significantly. The microcystin synthesis gene (*mcy*) showed a strong positive association (R² = 0.88) with cyanobacterial abundance. Moreover, cyanobacteria exhibited enhanced nutrient acquisition (*pstS* gene, R² = 0.69) and upregulated nitrogen metabolism pathways under warming conditions. Concurrently, we observed a marked increase in antibiotic resistance gene (ARG) abundance with rising temperatures. The relative abundance of multidrug resistance genes was consistently high across all conditions (50.82 % of total ARGs). ROS stress further intensified ARG proliferation, particularly for efflux pump genes (*e.g.*, *acrB*, *adeJ*, *ceoB*, *emrB*, *MexK*, *muxB*). Co-association network analysis identified key antibiotic-resistant bacteria (*e.g.*, *Streptococcus pneumoniae*, *Acinetobacter baylyi*) and ARGs (*e.g.*, *acrB*, *MexK*, *rpoB2*, *bacA*) central to resistance dissemination under warming conditions. This study demonstrates that even modest temperature increases (2-5°C) can promote both cyanobacterial blooms and antibiotic resistance in freshwater ecosystems over short time scales. The synergistic effects of temperature and oxidative stress underscore the complex challenges posed by climate change to water quality and public health.

**Highlights:**

- Short-term warming promotes toxic cyanobacteria and antibiotic resistance in freshwater.
- Harmful cyanobacteria and their metabolic potential increase under warming conditions.
- Reactive oxygen species mimic and exacerbate temperature effects.
- Oxidative stress response genes strongly correlate with ARG abundance.
- Co-association networks identify key host pathogen central to resistance spread.

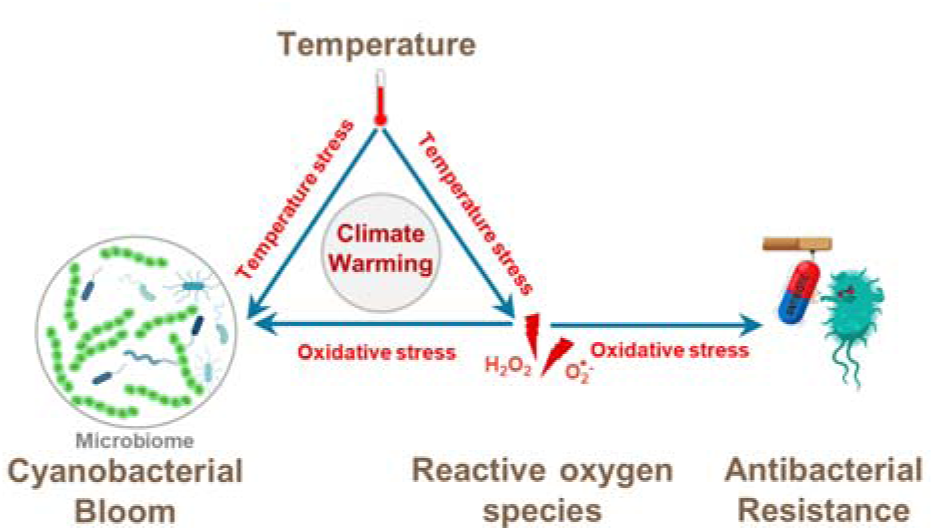

## 1. Introduction

Climate change poses an unprecedented threat to life on Earth, with far-reaching consequences for ecosystems, biodiversity, and human well-being (Cole et al., 2023; Sun et al., 2019). Climate warming has resulted in raising the average global temperature and more frequent high-temperature extremes. According to the sixth assessment report of the Intergovernmental Panel on Climate Change (IPCC), published in 2021, greenhouse gas emissions from anthropogenic activities have already warmed the climate by nearly 1.1°C since 1850-1900 (“Climate Change 2021—The Physical Science Basis,” 2021). The global average temperature is expected to reach or exceed 1.5°C within the next few decades (WMO, 2023). These changes will affect all regions of the Earth, including freshwater ecosystems, which provide vital services for aquatic life and human activities such as drinking water, food and recreation (Albert et al., 2021). Rising water temperatures are expected to cause a decrease in water quality, promoting eutrophication and the growth of algae, further exacerbating the problem of water scarcity (A. K et al., 2023). Moreover, warming conditions facilitate the formation of toxins, proliferation of antibacterial resistance genes (Li et al., 2022; Yang et al., 2023) (ARGs), waterborne pathogens, and an increase in the intensity and duration of algal and cyanobacterial blooms (O’Neil et al., 2012). Furthermore, both toxic cyanobacterial blooms and ARGs in freshwater are critical concerns for human health and well-being (Guo et al., 2018).

Cyanobacteria, among the oldest known photosynthetic organisms, first appeared on our planet roughly 3.5 billion years ago. (Bullerjahn and Post, 2014; Tomitani et al., 2006). The abundant growth of toxin-forming cyanobacteria in freshwater lakes, estuarine, and coastal ecosystems, due to the enrichment of certain inorganic and/or organic nutrients (nitrogen and phosphorus), has created serious concern worldwide (Conley et al., 2009; Fernández-González et al., 2022; Hernandez et al., 2021; Merel et al., 2013; Wood, 2016; Zhao et al., 2020). Studies have shown that nutrient over-enrichment in freshwater environments across three major European climate zones (Mediterranean, continental, and Atlantic) favors the growth of toxin-producing cyanobacteria (Filatova et al., 2020). Moreover, rising CO_2_ levels may result in a marked intensification of phytoplankton blooms in eutrophic and hypertrophic waters (Verspagen et al., 2014). Cyanobacteria play a key role in ecosystems as primary producers and generate various secondary metabolites, including harmful substances known as cyanotoxins. (Merel et al., 2013). Some common cyanotoxins and the cyanobacterial producer include microcystins (*e.g.*, *Anabaena*, *Arthrospira*, *Limnothrix*, *Microcystis*, *Nostoc*, *Oscillatoria* (*Planktothrix*), *Synechocystis*), nodularins (*Nodularia*), and saxitoxins (e.g., *Anabaena*, *Aphanizomenon*, *Planktothrix*, *Raphidiopsis*, and *Scytonema*) (Rastogi et al., 2015). Cyanotoxins can poison aquatic organisms, wildlife, pets, and humans through direct consumption of toxic cyanobacteria or ingestion of contaminated water (Rastogi et al., 2014). However, cyanotoxins may serve as a chemical defense, helping cyanobacteria outcompete other microorganisms and deter predators. (Amado and Monserrat, 2010; Berry et al., 2008; Jang et al., 2007). Recent studies have shown that metagenomics-based approaches hold great promise in identifying potential cyanotoxin- producing genera, such as *Microcystis*, *Planktothrix*, *Synechococcus*, *Cyanobium*, *Dolichospermum*, *Nodularia*, *Cylindrospermum*, and *Oscillatoria*, among others, in large rivers of the United States (Linz et al., 2023). Moreover, cyanobacteria have developed its defense mechanism against both temperature stress and oxidative stress including reactive oxygen species (ROS), which might enable it to thrive in a warming condition (Latifi et al., 2009; Yadav et al., 2022). As the emergence of a cyanobacterial bloom is the consequence of several coherent effects, including climate warming, nutrients, CO_2_ levels, and ability to mitigate oxidative stress, the impact of rising temperatures alone on these processes is poorly understood. In addition to cyanobacterial blooms, previous reports have indicated that climate warming could increase the threat of ARGs to environmental and human health (Qamar and Aatika, 2023). Metagenomics analysis of controlled temperature experiments revealed that increased temperature could remarkably reduce ARG diversity but increase ARG abundance, specifically for multidrug, tetracycline, and peptide resistance genes (Yu et al., 2023). Moreover, ARGs (*tetM*, *mecA*, *bacA*, *vatE*, and *tetW*) significantly increased with elevated water temperature, along with opportunistic pathogens like *Delftia*, *Legionella*, and *Pseudomonas*, implying that antibiotic resistance is a growing concern in a warming climate (Yu et al., 2023). As temperature stress has been linked with ROS stress (Fang et al., 2023; Hassan et al., 2020; Tignat-Perrier et al., 2022), the emergence of ARGs under warming temperatures could also be associated with ROS stress (Lu et al., 2018; Qi et al., 2023). We propose that elevated temperatures due to climate warming induce oxidative stress in aquatic microbial communities, leading to the proliferation of stress-tolerant, toxin-producing cyanobacteria and increased prevalence of ARGs. This occurs through both direct temperature stress on the microbiome and indirect effects *via* increased ROS. Cyanobacteria, being tolerant to both temperature and ROS stress, thrive under these conditions, while ROS stress simultaneously promotes the spread of ARGs. Thus, climate warming acts as a dual threat, fostering both cyanobacterial blooms and antibiotic resistance in aquatic ecosystems. In this study, we investigate impact of short-term temperature stress on microbes and the subsequent proliferation of stress-tolerant toxin-producing cyanobacteria and ARGs in Cosseys Reservoir, a crucial drinking water source in Auckland, New Zealand. By subjecting the lake’s microbial community to different temperature warming scenarios, simulating base (22°C), normal (24°C), and future (27°C) water temperatures, we aim to uncover the basis of temperature-induced proliferation of cyanobacteria and ARGs in freshwater lakes. Additionally, we performed explicit ROS addition to the temperature experiments to assess if oxidative stress could impact the microbial community in a manner similar to increased temperature stress. Using metagenomics analysis, our findings shed light on the potential impact of simulated short-term warming on freshwater ecosystems, particularly the susceptibility to cyanobacterial blooms and antibiotic resistance in the foreseeable future.

## 2. Materials and methods

### 2.1. Batch reactor experiments with different temperature and ROS levels

To investigate the proliferation of cyanobacterial blooms and ARGs under short-term temperature warming scenarios, batch reactor experiments were conducted using water samples from Cosseys Reservoir, Auckland, New Zealand. Three separate bioreactors with an effective volume of 1 L were used with different temperatures. Three temperature conditions were tested: a Baseline condition maintained at 22°C throughout, serving as a control for temperature effects; a Normal condition starting at 22°C and raised to 24°C; and a Future projected temperature condition starting at 22°C and raised to 27°C. All experiments were conducted over 48 hours, with systems kept for the first 24 hours to allow acclimatization of the microbial communities to the ambient temperature (22°C), then raised to their respective experimental temperatures for the remaining 24 hours. To assess the impact of ROS, exogenous addition of 75 µmol H_2_O_2_ was performed for two conditions: Base_H2O2_ (Baseline condition with added H_2_O_2_) and Normal_H2O2_ (Normal condition with added H_2_O_2_). The H2O2 range was selected based on our previous study. The purpose of these ROS-added controls was to compare whether ROS stress could mimic the effects of temperature stress on microbial systems.

### 2.2 DNA Extraction and Metagenomics Analysis

Samples of 100 mL were collected from reactors under different experimental conditions for DNA extraction. The extraction followed the protocol provided by the Qiagen® DNeasy PowerWater Kit (Qiagen, Germany), and the resultant DNA was preserved at -20°C for further analysis. The DNA samples were then sent off to the Auckland Genomics Centre (Auckland, NZ) to conduct metagenomic analysis. Because of the low concentration of DNA samples, a SpeedVac was used to concentrate the gDNA. A low input (<100 ng) protocol was then used to prepare libraries from 10-20 ng of total DNA. Libraries were made with the NEBNext® Ultra™ II DNA Library Prep Kit for Illumina® (E7645S) with unique dual indexes (cat no E7600). Amplification of the adapter-ligated DNA were performed via 9 cycles of PCR. Post library QC, normalization was carried out, followed by a final pool check using a bioanalyzer. The shotgun whole genome sequencing was carried out on a HiSeqX platform using 2x150 bp paired-end sequencing. The raw metagenome samples from these experiments can be accessed in the European Nucleotide Archive under the BioProject accession number PRJEB74089.

### 2.3 Bioinformatics Analysis

We performed preprocessing steps to eliminate low-quality sequences using Trimmomatic (Bolger et al., 2014) with specific quality thresholds: LEADING:3, TRAILING:3, SLIDINGWINDOW:10:15, and MINLEN:50 with the following settings: TruSeq3-PE- 2.fa:2:30:10:2:keepBothReads (Raes et al., 2021). For metagenomic taxonomic and functional profiling, we employed the SqueezeMeta v1.5.2 pipeline (Tamames and Puente-Sánchez, 2019). The read statistics per sample is provided in Table S1. Co-assembly was performed using Megahit (Li et al., 2015), and short contigs (<200 bps) were filtered out using prinseq (Schmieder and Edwards, 2011). Within the SqueezeMeta pipeline, Barrnap (Seemann, 2014) tool was used for predicting RNAs, while Prodigal (Hyatt et al., 2010) was utilized for predicting ORFs. Taxonomic ranks were assigned against NCBI GenBank nr database (Clark et al., 2016) with identify thresholds of 85, 60, 55, 50, 46, 42, and 40% for species, genus, family, order, class, phylum, and superkingdom ranks, respectively (Luo et al., 2014). Kyoto Encyclopedia of Genes and Genomes (Kanehisa and Goto, 2000), was used for functional assignments with Diamond (Buchfink et al., 2014) with a maximum e-value threshold of 1e-03 and sequence identify threshold of 50%. Moreover, the Comprehensive Antibiotic Resistance Database (Alcock et al., 2020) release 3.2.4 was used for annotating the ARGs and identifying the ARBs in metagenomics data. ARGs were annotated based on sequence identify of 70% and e-value threshold of 1e-03. ARGs conferring resistance to more than one antibiotics were classified as ‘multidrug resistant’ (MDR). Furthermore, classified species were cross-referenced with the known pathogens in the CARD database to identify ARBs in the metagenome. Bowtie2 (Langmead and Salzberg, 2012) was used for read mapping against the contigs. Both functional genes and ARGs were normalized to transcripts per million (TPM). The metagenomics data was analyzed using the SQMtools R package (Puente-Sánchez et al., 2020).

### 2.4 Statistical Analysis and Visualization

The co-association networks between the Cyanobacteria and other taxa and (ARGs and ARB pathogens) was constructed using the Sparse Correlations for Compositional data (SparCC) (Friedman and Alm, 2012) correlation method (Rho > 0.6 and P < 0.05) method in NetCoMi R package (Peschel et al., 2021). The co-association networks were visualized with Gephi (Bastian et al., 2009) version 1.10.1 using the Frucherman Reingold algorithm. Heatmaps were plotted to visualize changes in the abundance of cyanobacterial families and ARBs, utilizing a z-score with TBtools (Chen et al., 2020). The fold change analysis of pathways were preformed using Pathview R package (Luo and Brouwer, 2013). Correlation analysis between ARG and ROS stress response genes was conducted using OmicStudio tools (https://www.omicstudio.cn/). Graphics were generated using the ggplot2 (Wickham, 2011) package in R (R Core Team, 2022) version 4.2.1.

## 3. Results and Discussion

### 3.1. Temperature warming and oxidative stress restructure microbial communities and promote cyanobacterial proliferation

To investigate the impact of rising temperatures and oxidative stress on freshwater microbial communities, we subjected the microbiome of Cosseys Reservoir to three temperature scenarios: Base (22°C), Normal (24°C), and Future (27°C). Additionally, we explicitly introduced ROS (H_2_O_2_) to the Base and Normal treatments to evaluate the combined effects of temperature and oxidative stress. The initial microbial community (T0) was diverse, dominated by Proteobacteria (35.45%), Actinobacteria (10.28%), and Bacteroidetes (8.46%) (Figure 1a). Under baseline conditions (22°C), Proteobacteria increased significantly to 63.05%, while Actinobacteria and Bacteroidetes remained relatively stable. The addition of ROS at 22°C (Base_H2O2_) caused a marked decrease in Proteobacteria (36.42%) similar to T0, and an increase in Actinobacteria (18.46%) and Bacteroidetes (12.48%), suggesting that oxidative stress alters community structure. At the Normal temperature (24°C), Proteobacteria decreased (30.88%), while Actinobacteria (14.81%) and Bacteroidetes (11.94%) showed slight increases compared to the Base temperature. However, the addition of ROS at 24°C (Normal_H2O2_) dramatically increased Proteobacteria (57.49%), indicating a strong response to the combined temperature and ROS stressors. Under the Future temperature scenario (27°C), Proteobacteria (40.91%), Actinobacteria (17.07%), and Bacteroidetes (12.65%) were substantially present, highlighting significant community restructuring. Additionally, the initial cyanobacterial abundance was 6.11% (T0). At 22°C, it decreased slightly to 1.89%, but the addition of ROS to 22°C (Base_H2O2_) increased it to 13.39%, demonstrating the potential for oxidative stress to promote cyanobacterial growth. At 24°C, cyanobacterial abundance surged to 20.53%, indicating their sensitivity to temperature increase. The addition of ROS at 24°C (Normal_H2O2_) decreased cyanobacterial abundance to 7.86%, suggesting a complex combined effect of temperature and oxidative stress. Under the Future temperature scenario (27°C), cyanobacterial abundance reached 10.66%, significantly higher than the baseline but lower than the Normal condition. Bacteroidetes, Actinobacteria, and Proteobacteria have been previously reported to be associated with cyanobacterial bloom in freshwater pond (Xia et al., 2017). These findings suggest that rising temperatures alone can reshape freshwater microbial communities, promoting cyanobacterial proliferation during warmer climate. Previous short-term heat stress (29°C to 35°C) experiments also revealed shift in coral associated marine microbial communities (Ziegler et al., 2017). Moreover, these findings aligns with previous studies identifying temperature as a key factor in cyanobacterial growth (Chapra et al., 2017; Paerl and Paul, 2012). Notably, the cyanobacterial abundances observed in the Base_H2O2_ were similar to the Normal condition, and Normal_H2O2_ treatment were similar to Future temperature condition, implying that ROS stress could mimic the effects of temperature stress on cyanobacterial proliferation. These findings demonstrate that short-term temperature stress, exacerbated by oxidative stress, can significantly restructure freshwater microbial communities and promote the proliferation of cyanobacteria, increasing the risk of cyanobacterial blooms.

**Figure 1.**
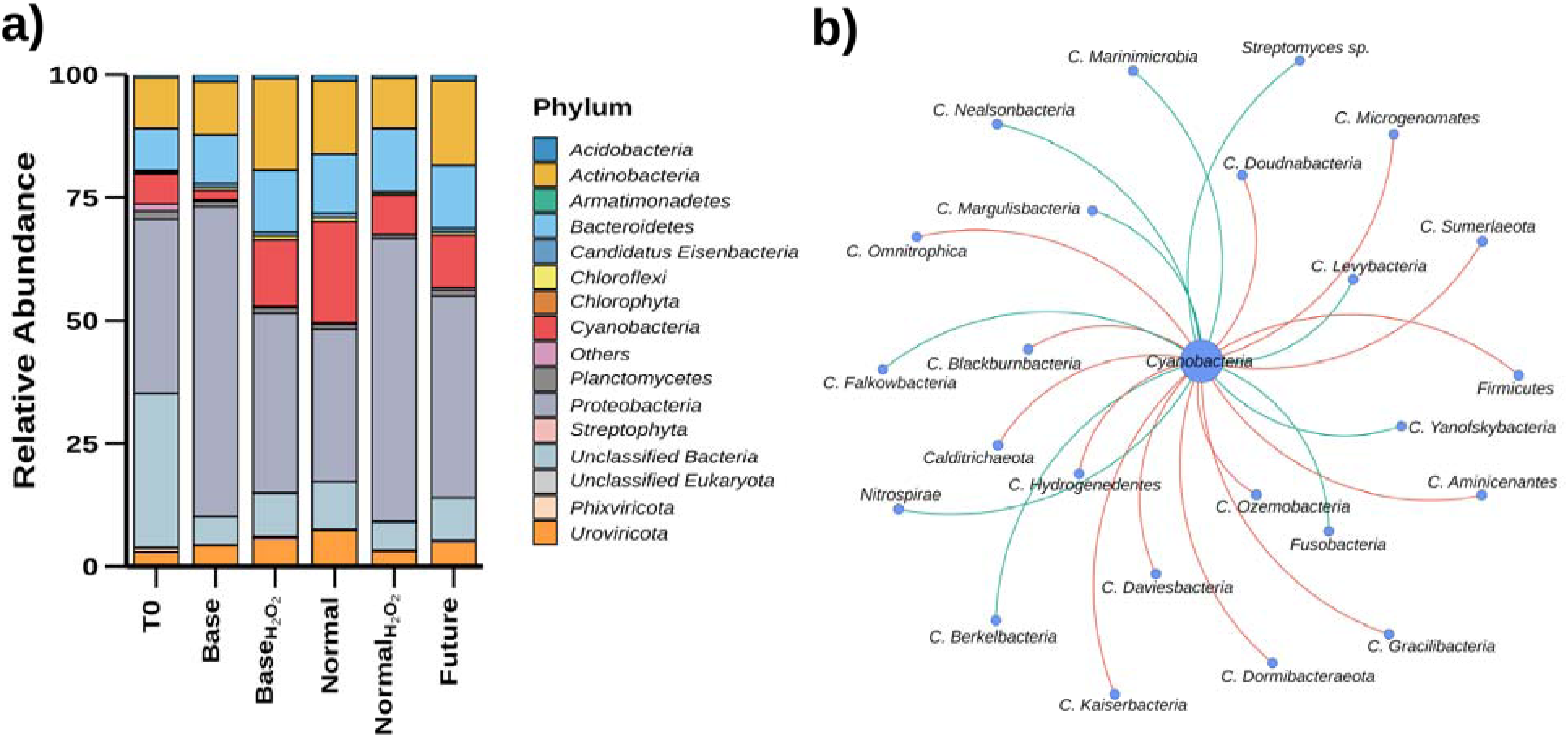
Changes in microbial community composition and cyanobacteria interactions under varying temperature and oxidative stress conditions. (a) Relative abundance of major microbial phyla in Cosseys Reservoir under different conditions: initial time point (T0), Baseline temperature (22°C), Base_H2O2_ (22°C + H_2_O_2_), Normal temperature (24°C), Normal_H2O2_ (24°C + ROS), and projected Future temperature (27°C). (b) Microbial co-association network highlighting interactions between cyanobacteria and other microbial taxa.

Furthermore, we constructed a co-association network between cyanobacteria and other phylum to understand the complex communication dynamics within the Cosseys Reservoir microbial community influencing the cyanobacterial bloom under increasing temperature conditions (Figure 1b, Table S2). The network analysis revealed intricate competitive and cooperative interactions involving cyanobacteria with 24 other taxa, out of which 19 belonged to the poorly characterized *Candidate* phylum. Cyanobacteria exhibited direct negative associations with nine taxa, including (*Calditrichaeota*, *Candidatus Dormibacteraeota*, *Candidatus Daviesbacteria*, *Candidatus Blackburnbacteria*, *Candidatus Kaiserbacteria*, *Candidatus Microgenomates*, *Candidatus Aminicenantes*, *Candidatus Omnitrophica*, *Candidatus Hydrogenedentes*, *Firmicutes*, *Candidatus Ozemobacteria*, *Candidatus Doudnabacteria*, *Candidatus Gracilibacteria*, and *Candidatus Sumerlaeota*), suggesting potential competition for resources. Notably, *Calditrichaeota* with the most negative association (Rho = -0.851) is a chemoorganoheterotrophic phylum, found in marine sediment (Marshall et al., 2017). Interestingly, genomic islands of *Caldithrix abyssi*, belonging to the novel bacterial phylum *Calditrichaeota* showed compositional and sequence similarities to genomic islands in cyanobacteria such as *Cyanothece*, *Anabaena*, *Nostoc* and *Spirosoma* (Kublanov et al., 2017). Moreover, *Firmicutes* (Rho = -0.687) are involved in decomposition and fermentation of organic matter in sediments, and its reduction in abundance has been observed during cyanobacterial blooms (Xia et al., 2017). These low abundant phyla with highly competitive relationships, likely influencing cyanobacterial population dynamics through resource competition. In contrast, cyanobacteria showed positive associations with ten taxa, including 7 *Candiate* phyla (*Candidatus Berkelbacteria*, *Candidatus Falkowbacteria*, *Candidatus Levybacteria*, *Candidatus Margulisbacteria*, *Candidatus Marinimicrobia*, *Candidatus Nealsonbacteria*, *Candidatus Yanofskybacteria*), *Nitrospirae*, *Streptomyces sp.*, and *Fusobacteria*, indicating potential cooperative interactions. The positive correlation suggests mutualistic relationships, such as nutrient exchange or metabolic support, underscoring the coexistence and mutualistic interactions within the community. These findings highlight the multifaceted nature of microbial interactions and their pivotal role in shaping cyanobacterial bloom dynamics in response to environmental changes like climate warming. While cyanobacteria face significant biotic pressure from competitive taxa, cooperative interactions with taxa like *Fusobacteria*, *Nitrospirae*, and a few *Candidate* phyla may enhance their resilience and adaptability. Understanding these complex relationships is crucial for predicting microbial community responses to future environmental changes and developing strategies to manage and mitigate the impacts of harmful cyanobacterial blooms on ecosystem health.

### 3.2 Proliferation of toxin-producing cyanobacterial families and functional adaptations under warming

We further analyzed the abundance of various cyanobacteria families to compare their relative changes across different treatment conditions (T0, Base, Base_H2O2_, Normal, Normal_H2O2_, Future). Most cyanobacteria families have higher relative abundance in the elevated temperature treatments (Normal and Future), indicating a marked increase in their relative abundance compared to the baseline and initial conditions (Figure 2a). Notably many of these abundant families are reported to by cyanotoxin producers (Jones et al., 2021). For instance, *Microcystaceae*, a well-known cyanotoxin (microcystin) producer, exhibited a notable increase in abundance from T0 to Normal and Future temperature, highlighting its proliferation under warming conditions. We also observed other cyanotoxin producer families including *Nostocaceae* (*g_Nostoc* and *g_Anabaena*), *Aphanizomenonaceae* (*g_Aphanizomenon*, *g_Dolichospermum*), *Leptolyngbyaceae* (*g_Leptolyngbya*), *Merismopediaceae* (*g_Synechocystis*), and *Microcoleaceae* (*g_Symploca*, *g_Planktothrix*) others. Notably, microcystins are prevalent in New Zealand’s lakes and reservoirs, with *Microcystis* being the primary producer of these toxins in planktonic samples (Puddick et al., 2019). Previous reports have also identified climate change will increase the severity, distribution and longevity of cyanobacterial blooms in lakes including the four most common pelagic bloom forming genera in New Zealand; *Aphanizomenon*, *Cylindrospermopsis*, *Dolichospermum* and *Microcystis* (Wood et al., 2017). Moreover, column clustering in the heatmap revealed that the treatments with elevated temperatures (Normal and Future) and those with oxidative stress (Base_H2O2_, Normal_H2O2_) cluster closely together. This indicates a similar response in cyanobacterial community composition under these conditions, suggesting that both temperature increase, and oxidative stress promote the proliferation of certain cyanobacteria families. Cyanobacteria, as oxygenic photosynthetic organisms, inevitably produce ROS during electron transport and developed a rapid antioxidant defenses for survival (Latifi et al., 2009). Regression analysis showed a strong association between SOD gene and cyanobacterial abundance (R² = 0.6415), while *perR* exhibited a weaker correlation (R² = 0.3736) (Figure S1). Both genes encode key oxidative defense enzymes in cyanobacteria (Latifi et al., 2009; Liu et al., 2023; Wang et al., 2011). Our findings reinforce the previous claims of climate warming induced proliferation of potentially toxigenic cyanobacteria (Chapra et al., 2017; Erratt et al., 2023; Paerl and Paul, 2012) and show the impact of temperature alone for promoting cyanobacterial blooms.

**Figure 2.**
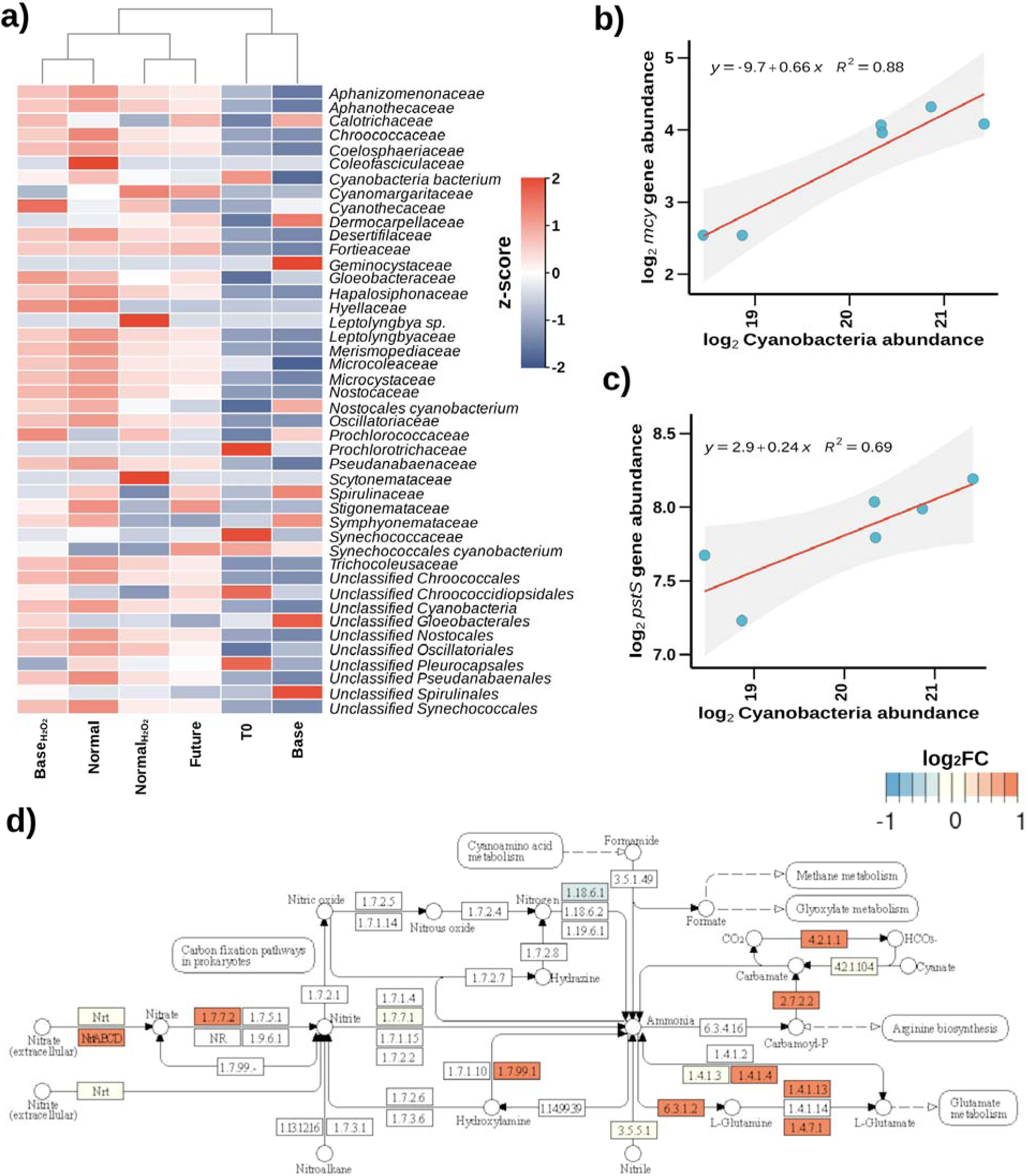
Cyanotoxin producers and functional potential of the cyanobacteria under different treatment conditions. (a) Heatmap showing the z-scores of relative abundances of different cyanobacteria families across various conditions. (b) Linear regression of *mcy* gene abundance (TPM) against total cyanobacterial abundance, showing a strong positive correlation (R² = 0.88), indicating increased toxin production potential with rising cyanobacterial populations. (c) Linear regression of *pstS* gene abundance against total cyanobacterial abundance, showing a positive correlation (R² = 0.69), suggesting enhanced nutrient acquisition capabilities with increasing cyanobacterial populations. (d) Nitrogen metabolism pathway highlighting the log_2_FC change (TPM) of different upregulated genes in the Future condition (27°C) compared to the Base condition (22°C).

Furthermore, the linear regression analysis shows a strong positive correlation (R² = 0.88) between the abundance of the *mcy* gene, responsible for microcystin production, and the total cyanobacterial abundance (Figure 2b). This indicated that cyanobacterial abundance is closely associated with increased *mcy* gene abundance, indicating a higher potential for toxin production under warming conditions. *mcy* gene has been previously used and an indicator of early detection of microcystin production (Duan et al., 2022; Padovan et al., 2023). Moreover, cyanobacterial metabolism has been associated with high nutrient uptake, specifically nitrogen and phosphate. We observed that the *pstS* gene, related to phosphate uptake, shows a positive correlation (R² = 0.69) with total cyanobacterial abundance (Figure 2c). As cyanobacterial populations grow, the expression of genes involved in phosphate acquisition also increased, suggesting an adaptation mechanism to optimize nutrient uptake in competitive environments. Additionally, the nitrogen metabolism was also upregulated in the future condition compared to the base condition (Figure 2d). The upregulation of key genes involved in nitrogen fixation, assimilation, and metabolism under future temperature scenarios indicates that cyanobacteria are not only proliferating but also actively altering their metabolic pathways to thrive in warmer environments. For example, genes involved in the uptake of extracellular nitrate (*nrtABCD*), transformation of nitrate to nitrite (*narB*) and subsequent assimilation into organic forms show higher metabolic potential, underscoring the adaptive responses of cyanobacteria to increased temperatures. Moreover, cyanobacterial photosynthesis and carbon-fixation related genes were also present in higher abundance in the Future temperature compared to the Base temperature (Figure S2 and Figure S3). The results clearly demonstrate that rising temperatures and oxidative stress significantly promote the proliferation of cyanotoxin-producing cyanobacteria. The strong correlations between cyanobacterial abundance and the presence of *mcy* and *pstS* genes indicate an increased potential for toxin production and efficient nutrient uptake. Furthermore, the increased potential of nitrogen metabolism and photosynthesis pathways in warmer conditions underscores the adaptive mechanisms that enable cyanobacteria to dominate in changing environmental conditions.

### 3.3 Temperature and ROS induced proliferation of antibacterial resistance

In addition to the effects on cyanobacterial proliferation, we further assessed the relative abundance of various ARGs under different treatment conditions to understand the impact of increased temperature and oxidative stress. In total, 122 ARGs were detected in the metagenomic assembly, which were further classified into 45 families, 20 drug classes and 8 different mechanisms of resistance based on the CARD database (Alcock et al., 2020). The most common ARG families were resistance-nodulation-cell division (RND) antibiotic efflux pump (32.79 %), major facilitator superfamily (MFS) antibiotic efflux pump (8.20 %), miscellaneous ABC-F subfamily ATP-binding cassette (ABC) antibiotic efflux pump (5.74 %), and tetracycline- resistant ribosomal protection protein (5.74 %). In terms of mechanism of resistance, antibiotic efflux (46.72 %), antibiotic inactivation (22.13 %), antibiotic target protection (17.21 %), antibiotic target alteration (7.38 %) and antibiotic target replacement (3.28 %) were most prevalent. The analysis of ARG profiles across different conditions revealed dynamic changes in resistance mechanisms. The initial T0 condition showed low relative abundance of ARGs, representing a baseline level of antibiotic resistance in the Cosseys Reservoir microbial community. With increasing temperature, total ARG abundance increased substantially in Base, Normal, and Future conditions compared to T0. Notably, the highest ARG abundance was observed under oxidative stress conditions (Base_H2O2_ and Normal_H2O2_), suggesting a strong link between oxidative stress and the proliferation of ARGs (Figure 3a). Several antibiotic ARGs showed consistent presence, across all experimental conditions, indicating their fundamental role in baseline resistance mechanisms. For example, the rifamycin-resistant beta-subunit of RNA polymerase (*rpoB2* and *Bado_rpoB_RIF*) was observed along with the peptide antibiotic- resistant *ugd* gene, and multidrug-resistant RND efflux pumps (*MexK* and *MuxB*), highlighting the mechanisms in baseline resistance. Moreover, the baseline resistance in T0 predominantly involved antibiotic target alteration/protection/inactivation, and efflux systems, with genes like *rpoB2*, *ceoB*, *Bado_rpoB_RIF*, *tet(36)*, *ugd*, *MuxB*, *TEM-116*, and *MexK*. Under the Base condition, a shift occurred towards additional resistance through cell wall synthesis (*bacA*) and efflux pumps (*mtrA*), with genes such as *vatF* and *SFH-1* becoming more abundant. The presence of oxidative stress in both Base_H2O2_ and Normal_H2O2_ conditions further intensified efflux and stress response mechanisms, as evidenced by the increased presence of *oleC*, *MuxC*, *arnA*, and *emrB*. The Normal condition highlighted tetracycline resistance (*tet(36)*, *tetB(P)*), efflux pumps (*MuxB*, *mdtB*, *rsmA*), while the Future condition underscores a combination of efflux (*oleC*, *MuxC*) and adaptive regulatory mechanisms. Moreover, 9 ARGs showed an increasing trend with rising temperature, out of which six were antibiotic efflux pumps (*AcrF*, *lmrC*, *mdsB*, *oleC*, *poxtA*, *RanA*), and three involved with antibiotic targets (*otr(A)S.liv*, *rpoB2*, *vgaC*). These findings demonstrate that antibiotic resistance is highly adaptable, with specific ARGs being abundant in response to temperature and ROS stress, suggesting that future warming scenarios could sustain high levels of antibiotic resistance in aquatic microbial communities in agreement with previous report (Yu et al., 2023).

**Figure 3.**
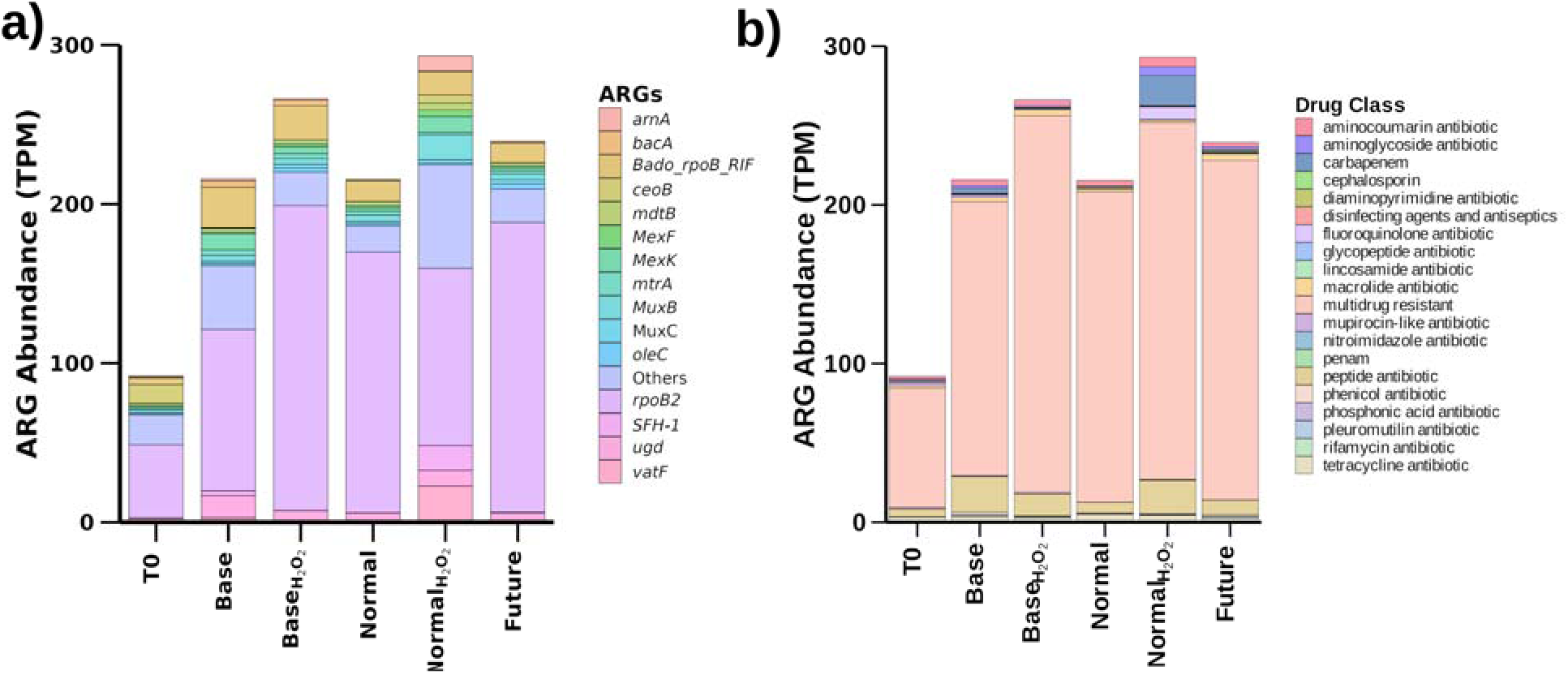
Stacked bar chart showing the relative abundance of different a) ARGs and b) drug classes under various treatment conditions, highlighting the enhanced antibiotic resistance potential in response to temperature and ROS.

Furthermore, the analysis of antibiotic resistance patterns across different drug classes revealed several key trends (Figure 3b). Key resistance drug classes identified in the total metagenome assembly included MDR (50.82 %), tetracycline antibiotic (8.20 %), and peptide antibiotic (6.56 %), aminoglycoside antibiotic (4.10 %), carbapenem (4.10 %) and macrolide antibiotic (3.28 %). Notably, the relative abundance of MDR genes was consistently high across all conditions, suggesting a pervasive and exacerbated MDR resistance under environmental stressors. Additionally, resistance to aminocoumarin, aminoglycoside, and carbapenem antibiotics showed significant increases under the combined effects of higher temperatures and oxidative stress, indicating the potential for these stressors to enhance resistance to these drug classes. Peptide antibiotic resistance also increased under environmental stress conditions, highlighting the impact of environmental changes on resistance to this drug class. However, tetracycline resistance exhibited a variable response, with higher abundance under normal conditions and a slight decrease under future conditions. In agreement with these findings, multidrug, tetracycline and peptide resistance genes had exhibited significant increase under warming temperature (Yu et al., 2023).

Rising temperatures significantly increased ARG abundance, consistent with previous warming experiments (Li et al., 2022; Yu et al., 2023). However, our results indicated that ROS alone could influence ARG enrichment, leading to further analysis of correlations between ROS stress response genes and ARGs under increasing temperatures (Figure S4). We examined correlations between ARGs and ROS stressor genes (*oxyR*: hydrogen peroxide-inducible genes activator, *soxR*: redox-sensitive transcriptional activator, *SOD*: superoxide dismutase, *katG*: catalase-peroxidase, *perR*: peroxide stress response regulator) during temperature treatment, identifying 25 unique ARGs correlated to different ROS stressors. The strongest associations were with *oxyR* (7 ARGs), *soxR* (18 ARGs), and *SOD_Cu-Zn_* (17 ARGs). Prevalent ARGs across conditions (*rpoB2*, *Bado_rpoB_RIF*, *MexK*, *ugd*) showed strong correlation with *oxyR*. Notably, several ARGs were shared across multiple ROS stressors, suggesting an overlap between oxidative stress response and antibiotic resistance. The most shared ARGs were *ceoB* and *tet(W)*, correlated with five and four ROS stressors, respectively. Other frequently shared ARGs included *rpoB2*, *MexK*, *ugd*, and *acrB*. This overlap indicated interconnected bacterial responses to oxidative stress and antibiotic pressure. The consistent presence of efflux pump genes (*e.g.*, *MexK*, *acrB*) across multiple ROS stressors suggests a dual role in expelling both antibiotics and oxidative agents. Notably, 15 of these ARGs exhibited antibiotic resistance through efflux pumps, indicating a strong association between ROS stress and this resistance mechanism. ROS stress has been previously linked with ARGs (Ma et al., 2021), specifically efflux pumps (Jin et al., 2018; Li et al., 2021; Qi et al., 2023).

Moreover, the abundance of ARB exhibited distinct patterns across the experimental conditions, with some ARBs proliferating under simulated future climate scenarios while others experienced a decline (Figure S5). The total relative abundance of ARBs decreased significantly under experimental conditions compared to the initial reservoir sample (T0). However, certain ARBs like *Caulobacter vibrioides*, *Chryseobacterium indologenes*, and *Citrobacter braakii* increased considerably in simulated future conditions. *Elizabethkingia meningoseptica* and *Stenotrophomonas maltophilia* also showed marked increases, indicating potential resilience. *Pseudomonas fluorescens* and *Caulobacter vibrioides* consistently increased in future conditions despite negative responses in other treatments. *Burkholderia pseudomallei* and *Pantoea agglomerans* demonstrated significant increase, underscoring their adaptability. Responses to ROS treatment varied, with *Legionella pneumophila* responding positively and *Aeromonas hydrophila* negatively, highlighting differential resistance mechanisms among ARBs.

We further employed a co-association network approach to analyze the relationships between ARB and ARG under increasing temperatures (Lira et al., 2020). Using the SparCC correlation method (Rho > 0.6 and P < 0.05), we generated a cumulative network and identified key ARBs and ARGs based on their degree centrality measures (Figure 4 and Table S3). *Streptococcus pneumoniae*, *Acinetobacter baylyi*, and *Aeromonas veronii* emerged as the most central ARBs, connected to a large number of ARGs (43, 41, and 40, respectively), underscoring their significant role as reservoirs of antibiotic resistance. Other notable ARBs with high degree centrality included *Aeromonas jandaei*, *Acinetobacter baumannii*, *Acinetobacter venetianus*, *Acinetobacter johnsonii*, *Acinetobacter haemolyticus*, *Vibrio cholerae*, and *Klebsiella pneumoniae*, associated with numerous ARGs. Additionally, the ARGs *acrB*, *MexK*, *rosB*, *rpoB2*, *bacA*, *adeB*, *abeM*, *ugd*, *poxtA*, *Bado_rpoB_RIF*, *adeI*, *MuxB*, *adeC*, *adeJ*, *adeR*, and *adeA* were identified as highly central in the network, linked to multiple ARBs, indicating their widespread presence and potential for horizontal gene transfer during warming temperature (MacFadden et al., 2018). The high connectivity of certain ARBs and ARGs suggests their crucial roles in maintaining the network and the potential for rapid dissemination of resistance genes. The identification of these central nodes provides valuable targets for intervention strategies, as disrupting them may mitigate the spread of antibiotic resistance. The findings underscore the importance of understanding the network dynamics of ARBs and ARGs in the context of temperature stress. As temperatures increase, the interaction network between ARBs and ARGs may become more pronounced, potentially accelerating the spread of resistance across different bacterial populations.

**Figure 4.**
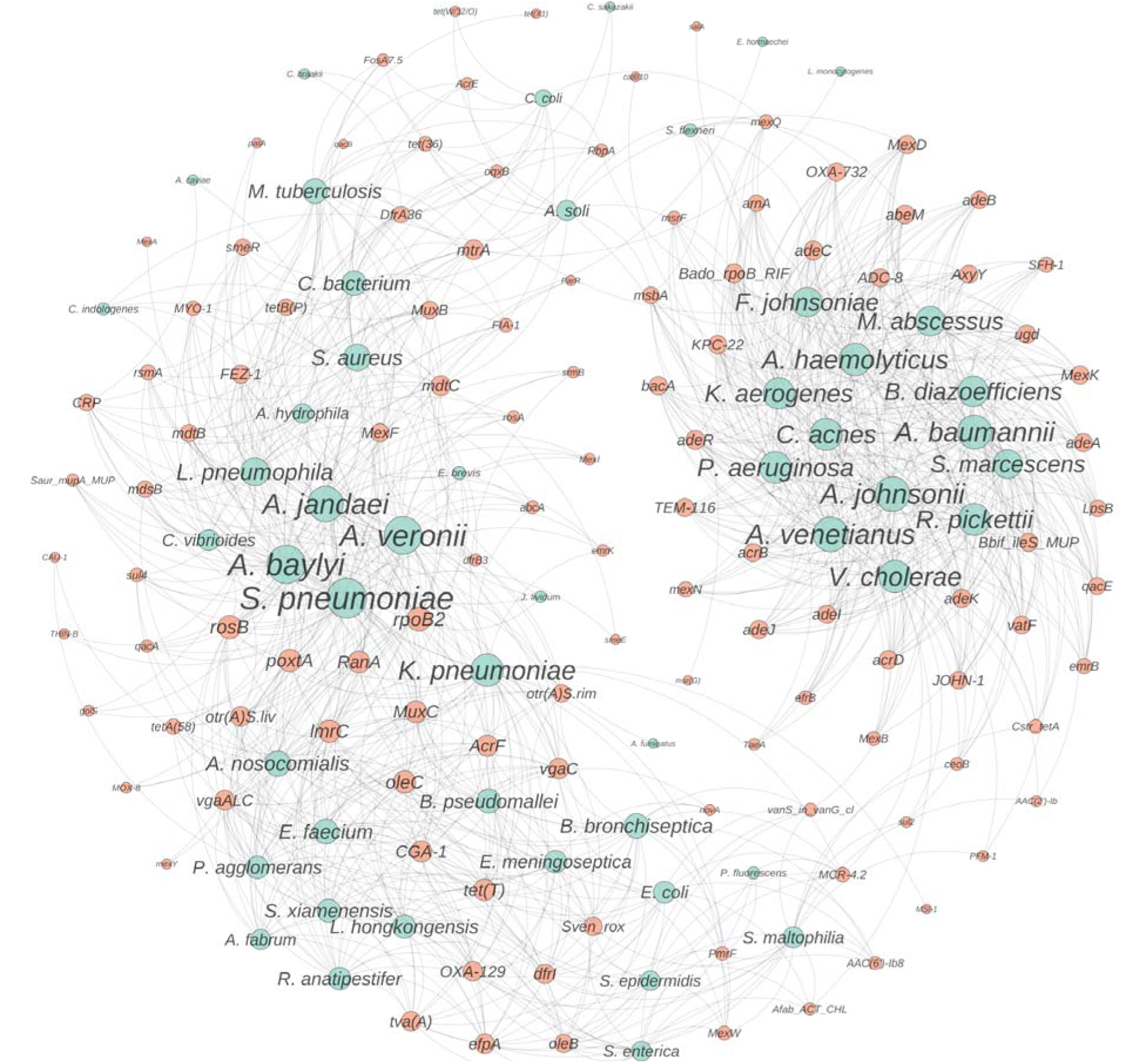
Co-association network of ARBs and ARGs. The network illustrates the interactions between ARBs (blue nodes) and ARGs (orange nodes) under temperature warming conditions. Node size represents the degree centrality, indicating the number of connections a node has within the network. Larger nodes have higher centrality, signifying their critical role in the network. Edges represent co-association relationships between ARBs and ARGs.

## 4. Conclusion

This study provides novel insights into the dual threats of toxic cyanobacterial blooms and antibiotic resistance proliferation in freshwater ecosystems under short-term climate warming scenarios. Our innovative approach, combining temperature manipulation with oxidative stress induction, revealed that even modest temperature increases (2-5°C) can significantly alter microbial community structures over short time scales. Harmful cyanobacterial species, particularly *Microcystaceae*, along with enhanced cyanobacterial metabolic potential under warming conditions. Concurrently, we observed a concerning rise in ARG abundance, strongly correlated with oxidative stress response genes. However, these findings are limited to metagenomic potential and need to be further validated by future metatranscriptomics or metaproteomic expression-based confirmatory studies. Notably, our study shows that reactive

oxygen species can mimic and exacerbate temperature effects on freshwater microbial communities, providing a potential mechanistic link between temperature stress and microbial community changes. While the short-term nature of our experiments and focus on a single reservoir limit generalizability, our findings underscore the urgent need for proactive measures to mitigate climate change effects and develop strategies to manage cyanobacterial blooms and combat antibiotic resistance in aquatic environments. This research advances our understanding of how climate warming may reshape freshwater microbial communities, with far-reaching implications for ecosystem health, water quality, and public health, providing a foundation for future long-term studies across diverse freshwater ecosystems.

## Associated content Supporting Information

All the supplementary tables and figures are provided in Supporting Information.

## ORCID

Bharat Manna: 0000-0001-7394-8204

Naresh Singhal: 0000-0002-9461-688X Wensi Zhang: 0000-0002-5515-2306

## Author Contributions

BM: Conceptualization, Methodology, Formal analysis, Visualization, Writing - original draft. EJ: Conceptualization, Methodology. Writing - review & editing. WZ: Methodology, Writing - review & editing. XZ: Methodology, Writing - review & editing. BL: Methodology, Writing - review & editing. GMT: Methodology, Writing - review & editing. NS: Conceptualization, Funding acquisition, Supervision, Writing - review & editing.

## Notes

The authors declare no competing financial interest.

## Acknowledgment

The authors acknowledge the use of New Zealand eScience Infrastructure (NeSI) high performance computing facilities, consulting support and/or training services as part of this research. New Zealand’s national facilities are provided by NeSI and funded jointly by NeSI’s collaborator institutions and through the Ministry of Business, Innovation & Employment’s Research Infrastructure programme. URL https://www.nesi.org.nz. The authors also acknowledge the Centre for eResearch at the University of Auckland for their help in facilitating this research. http://www.eresearch.auckland.ac.nz.

