## Supporting Information for "Short-Term Warming Induces Cyanobacterial Blooms and Antibiotic Resistance in a Freshwater Lake as Revealed by Metagenomics Analysis"

**for**

Table of Contents Pages

Tables S1-S3, pages S2-S5

Figures S1-S5, pages S6-S10

**Table S1. Details on the read statistics from the metagenomics assembly.**

| Sample | Number of Reads | Number of ORFs | Number of KEGG Annotation | Number of CARD Annotation |
|---|---|---|---|---|
| **T0** | 32009332 | 1235997 | 413958 | 297 |
| **Base** | 42706716 | 1934764 | 773522 | 605 |
| **Base$_{H2O2}$** | 40789728 | 1972225 | 743212 | 565 |
| **Normal** | 32665968 | 1745506 | 651586 | 463 |
| **Normal$_{H2O2}$** | 37098554 | 1800392 | 695723 | 572 |
| **Future** | 34178138 | 1848123 | 688499 | 496 |

**Table S2. Co-association between cyanobacteria and other taxa (Rho cut-off 0.6 and P < 0.05).**

| Interacting Taxa | Association Values | Interaction Type |
|---|---|---|
| *Calditrichaeota* | -0.851057392 | Negative |
| *Candidatus Aminicenantes* | -0.741259933 | Negative |
| *Candidatus Berkelbacteria* | 0.723965827 | Positive |
| *Candidatus Blackburnbacteria* | -0.774548466 | Negative |
| *Candidatus Daviesbacteria* | -0.803798674 | Negative |
| *Candidatus Dormibacteraeota* | -0.82908773 | Negative |
| *Candidatus Doudnabacteria* | -0.667298643 | Negative |
| *Candidatus Falkowbacteria* | 0.81002238 | Positive |
| *Candidatus Gracilibacteria* | -0.664900112 | Negative |
| *Candidatus Hydrogenedentes* | -0.705752705 | Negative |
| *Candidatus Kaiserbacteria* | -0.760803655 | Negative |
| *Candidatus Levybacteria* | 0.622565031 | Positive |
| *Candidatus Margulisbacteria* | 0.856224294 | Positive |
| *Candidatus Marinimicrobia* | 0.630504913 | Positive |
| *Candidatus Microgenomates* | -0.741756729 | Negative |
| *Candidatus Nealsonbacteria* | 0.770185423 | Positive |
| *Candidatus Omnitrophica* | -0.707813088 | Negative |
| *Candidatus Ozemobacteria* | -0.667737977 | Negative |
| *Candidatus Sumerlaeota* | -0.609168143 | Negative |
| *Candidatus Yanofskybacteria* | 0.806397463 | Positive |
| *Firmicutes* | -0.687736073 | Negative |
| *Fusobacteria* | 0.880943897 | Positive |
| *Nitrospirae* | 0.644060638 | Positive |
| *Streptomyces sp.* | 0.666546614 | Positive |

**Table S3. List of abbreviations used for ARGs node labels**

| Name | Node label |
|---|---|
| *Acinetobacter baumannii* | *A. baumannii* |
| *Acinetobacter baylyi* | *A. baylyi* |
| *Acinetobacter haemolyticus* | *A. haemolyticus* |
| *Acinetobacter johnsonii* | *A. johnsonii* |
| *Acinetobacter nosocomialis* | *A. nosocomialis* |
| *Acinetobacter soli* | *A. soli* |
| *Acinetobacter venetianus* | *A. venetianus* |
| *Aeromonas caviae* | *A. caviae* |
| *Aeromonas hydrophila* | *A. hydrophila* |
| *Aeromonas jandaei* | *A. jandaei* |
| *Aeromonas veronii* | *A. veronii* |
| *Agrobacterium fabrum* | *A. fabrum* |
| *Aspergillus fumigatus* | *A. fumigatus* |
| *Bordetella bronchiseptica* | *B. bronchiseptica* |
| *Bradyrhizobium diazoefficiens* | *B. diazoefficiens* |
| *Burkholderia pseudomallei* | *B. pseudomallei* |
| *Campylobacter coli* | *C. coli* |
| *Caulobacter vibrioides* | *C. vibrioides* |
| *Chryseobacterium indologenes* | *C. indologenes* |
| *Citrobacter braakii* | *C. braakii* |
| *Clostridiaceae bacterium* | *C. bacterium* |
| *Cronobacter sakazakii* | *C. sakazakii* |
| *Cutibacterium acnes* | *C. acnes* |
| *Elizabethkingia meningoseptica* | *E. meningoseptica* |
| *Empedobacter brevis* | *E. brevis* |
| *Enterobacter hormaechei* | *E. hormaechei* |
| *Enterococcus faecium* | *E. faecium* |
| *Escherichia coli* | *E. coli* |
| *Flavobacterium johnsoniae* | *F. johnsoniae* |
| *Janthinobacterium lividum* | *J. lividum* |
| *Klebsiella aerogenes* | *K. aerogenes* |
| *Klebsiella pneumoniae* | *K. pneumoniae* |
| *Laribacter hongkongensis* | *L. hongkongensis* |
| *Legionella pneumophila* | *L. pneumophila* |
| *Listeria monocytogenes* | *L. monocytogenes* |
| *Mycobacterium tuberculosis* | *M. tuberculosis* |
| *Mycobacteroides abscessus* | *M. abscessus* |
| *Pantoea agglomerans* | *P. agglomerans* |
| *Pseudomonas aeruginosa* | *P. aeruginosa* |
| *Pseudomonas fluorescens* | *P. fluorescens* |
| *Ralstonia pickettii* | *R. pickettii* |
| *Riemerella anatipestifer* | *R. anatipestifer* |
| *Salmonella enterica* | *S. enterica* |
| *Serratia marcescens* | *S. marcescens* |
| *Shewanella xiamenensis* | *S. xiamenensis* |
| *Shigella flexneri* | *S. flexneri* |

| | |
|---|---|
| *Staphylococcus aureus* | *S. aureus* |
| *Staphylococcus epidermidis* | *S. epidermidis* |
| *Stenotrophomonas maltophilia* | *S. maltophilia* |
| *Streptococcus pneumoniae* | *S. pneumoniae* |
| *Vibrio cholerae* | *V. cholerae* |

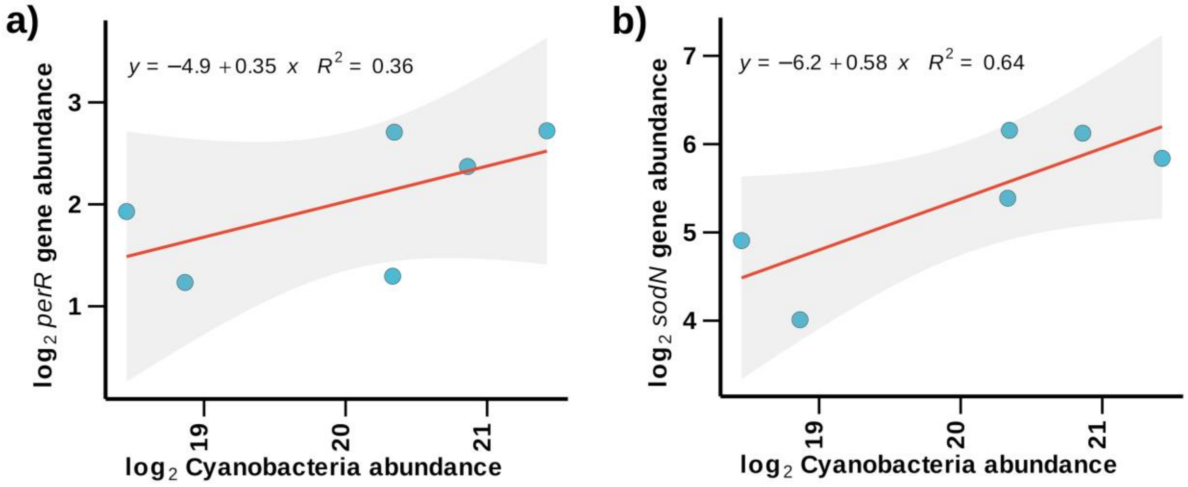


**Figure S1.** Linear regression of (a) *perR* (peroxide stress response regulator) and (b) *sodN* (nickel superoxide dismutase) gene abundance against total cyanobacterial abundance, showing a positive association, indicating a ROS stress response in cyanobacterial populations.

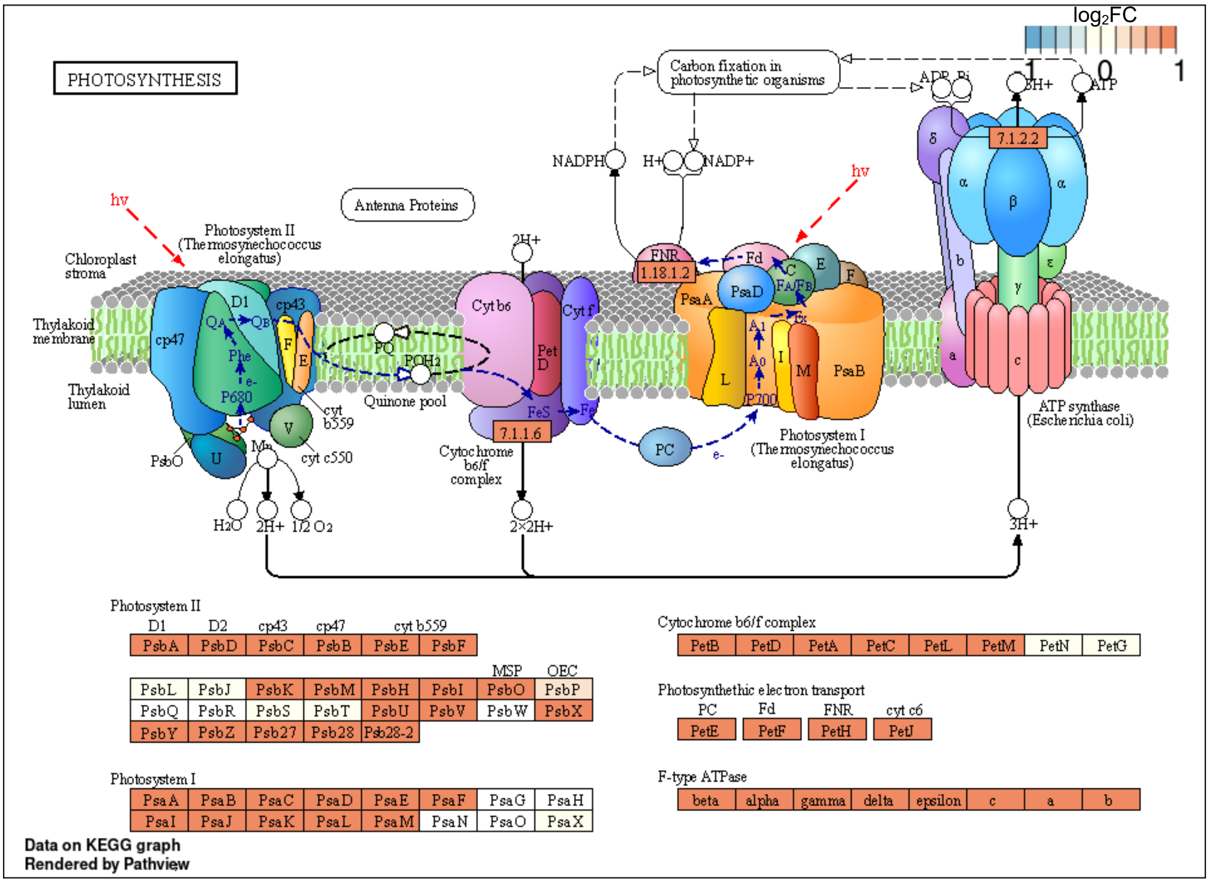


**Figure S2.** Photosynthesis pathway highlighting the log$_2$FC change (TPM) of multiple upregulated genes in the Future condition (27°C) compared to the Base condition (22°C).

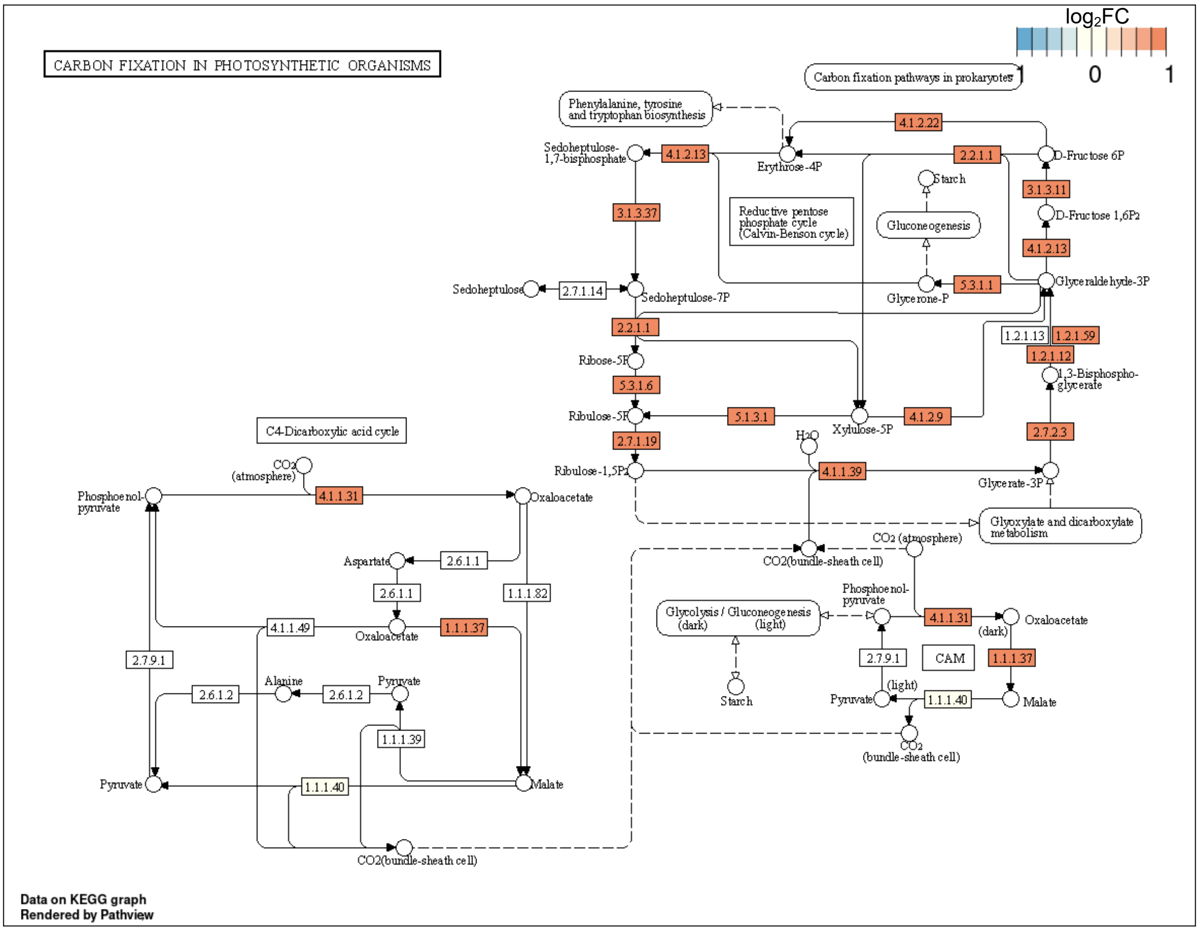


**Figure S3.** Carbon fixation in photosynthetic organisms highlighting the $log_2FC$ change (TPM) of multiple upregulated genes in the Future condition (27°C) compared to the Base condition (22°C).

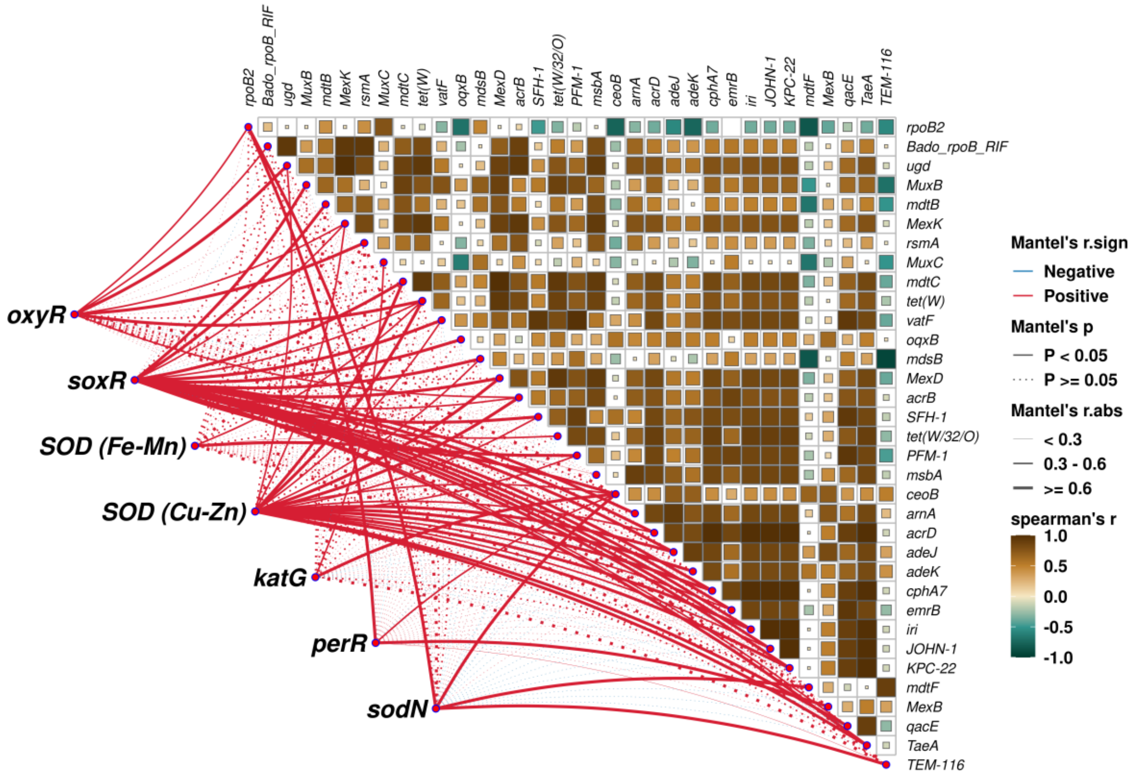


**Figure S4.** A network analysis depicting the correlations between selected ARGs and ROS stress response genes during the increasing temperature treatment. *oxyR*: hydrogen peroxide-inducible genes activator, *soxR*: redox-sensitive transcriptional activator, *SOD*: superoxide dismutase, *katG*: catalase-peroxidase, *perR*: peroxide stress response regulator, *sodN*: nickel superoxide dismutase.

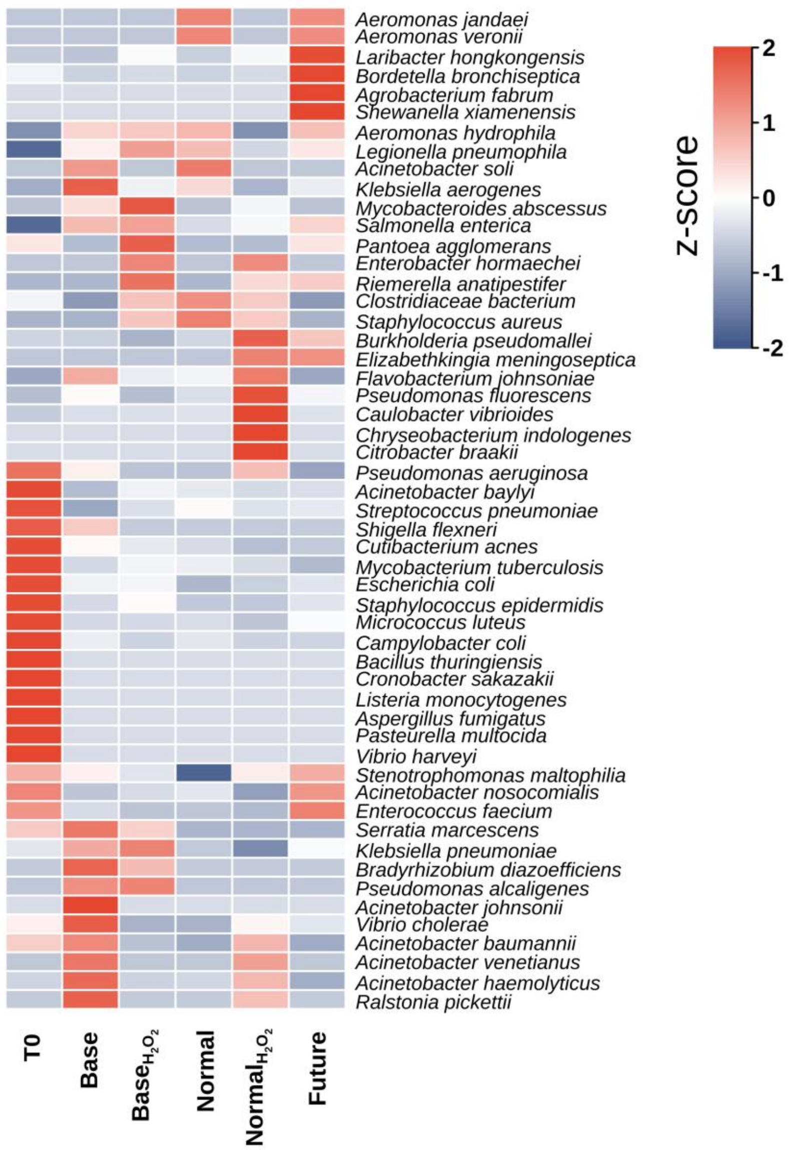


**Figure S5.** Heatmap showing the z-scores of relative abundances of different antibiotic resistant bacteria (ARBs) across various conditions. Species identified in the metagenomics analysis were searched against the list of known pathogens in CARD database to identify the ARBs.
